# Memorize-Generalize: An online algorithm for learning higher-order sequential structure with cloned Hidden Markov Models

**DOI:** 10.1101/764456

**Authors:** Rajeev V. Rikhye, J. Swaroop Guntupalli, Nishad Gothoskar, Miguel Lázaro-Gredilla, Dileep George

## Abstract

Sequence learning is a vital cognitive function and has been observed in numerous brain areas. Discovering the algorithms underlying sequence learning has been a major endeavour in both neuroscience and machine learning. In earlier work we showed that by constraining the sparsity of the emission matrix of a Hidden Markov Model (HMM) in a biologically-plausible manner we are able to efficiently learn higher-order temporal dependencies and recognize contexts in noisy signals. The central basis of our model, referred to as the Cloned HMM (CHMM), is the observation that cortical neurons sharing the same receptive field properties can learn to represent unique incidences of bottom-up information within different temporal contexts. CHMMs can efficiently learn higher-order temporal dependencies, recognize long-range contexts and, unlike recurrent neural networks, are able to natively handle uncertainty. In this paper we introduce a biologically plausible CHMM learning algorithm, memorize-generalize, that can rapidly memorize sequences as they are encountered, and gradually generalize as more data is accumulated. We demonstrate that CHMMs trained with the memorize-generalize algorithm can model long-range structure in bird songs with only a slight degradation in performance compared to expectation-maximization, while still outperforming other representations.

## Introduction

The ability to identify and process temporal sequences within complex sensory signals is an important cognitive function. For example, during speech, our interpretation of sentences depends on our ability to integrate words and their embedded contexts over time (Doupe & Kuhl, 2002). In this paper, we describe a biologically plausible sequence learning algorithm —memorize-generalize on cloned HMMs— and demonstrate its ability to discover higher order temporal structure in bird-song sequences.

Understanding how the brain represents temporal sequences has been a major effort in neuroscience (Clegg, DiGirolamo, & Keele, 1998; Abeles, 1991). In general, the cortex is able to learn continuously from noisy and incomplete data, and excels at making probabilistic predictions of future events based on current context (Rao & Ballard, 1999). As such, an intelligent sequence learning algorithm should ideally be able to learn online, be robust to noisy data, and must be able to make higher-order contextual predictions (Cui, Ahmad, & Hawkins, 2016).

Cloned HMMs (CHMM) were introduced in our earlier work, and shown to be capable of modeling higher order sequences (Dedieu et al., 2019). Unlike a traditional HMM which has a dense emission matrix, the CHMM has a particular sparsity structure where multiple hidden states, or clones, are deterministically mapped to the same observation. This sparsity structure is inspired by the connectivity observed in the cortex, where, depending on their intra-columnar connectivity, different neurons with the same bottom-up receptive field can be used to encode different sequential contexts (Hawkins, George, & Niemasik, 2009). By constraining the sparsity of the emission matrix, the CHMM outperforms Long Short Term Memory (LSTMs) networks and n-grams at learning higher-order temporal structure (Dedieu et al., 2019), and is robust to uncertainty in the temporal sequence.

While CHMMs were able to learn higher-order sequences, the expectation-maximization (EM) algorithm we used to train the model required numerous repetitions of the sequences to learn them, and strict memorization of the sequences might not even occur. In biological sequence learning, it is advantageous to be able to rapidly memorize recent experience and then gradually generalize that with repetitions or with accumulation of more data. Memorize-generalize, which we introduce in the following section, is such an algorithm.

The rest of this paper is organized as follows: We first describe the CHMM model, the properties of its representational structure and the biological inspiration behind it. Next, we describe the memorize-generalize learning algorithm for CHMMs. Finally, we demonstrate the performance of this algorithm in modeling bird songs and compare to CHMMs trained with EM.

## Cloned Hidden Markov Model

The CHMM, which we describe in detail in (Dedieu et al., 2019), is a cortically-inspired sequence learning algorithm that meets all three of the above mentioned criteria. The primary difference between the CHMM and the HMM is the unique sparsity structure of the emission matrix that deterministically maps multiple hidden states (clones) to the same emission state (Fig. 1A). These clones emit the same observed symbol, and through learning of the transition matrix, different clones represent different temporal contexts in which the symbol occurs in the training data. Even though having a deterministic, sparse, emission matrix makes the CHMM less expressive compared to an HMM (Dedieu et al., 2019), this property makes training CHMMs via EM less susceptible to finding poor local minima.

**Figure 1:**
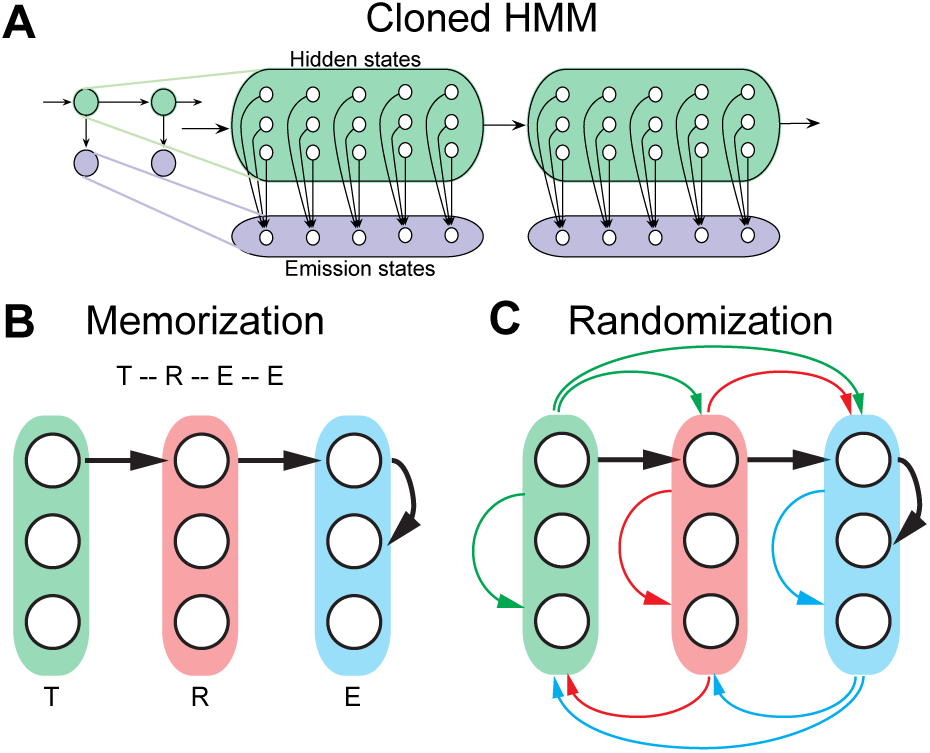
Illustration of the memorize-generalize algorithm. (A) Illustration of a CHMM. (B) Consider a CHMM with 3 clones learning the sequence T-R-E-E. Each colored column represents clones for each character. During the memorization, the CHMM represents the transition between each character as shown by the black arrows. (C) Endowing the CHMM with a random jump probability is akin to adding an all-to-all fully connected transition matrix. Note, the colored arrows indicate all-to-all connectivity between the clones for each character.

Our rationale for using this particular sparsity structure is based on the following observations. First, each cortical column contains several neurons that share similar bottom-up receptive fields (Hawkins et al., 2009). For example, neighboring neurons in Layer 2/3 derive their receptive fields from common feed-forward input from neurons in Layer 4 and thus would redundantly encode the same stimulus features (Yen, Baker, & Gray, 2007). Second, although these neurons share a common receptive field, they respond to different instances of the same sensory information (Vinje & Gallant, 2000). As a result, different neurons can learn to represent unique occurrences of information in different sequences.

The idea of cloning the states of a Markov chain to create higher-order representations was first proposed in a popular compression algorithm, dynamic Markov coding (Cormack & Horspool, 1987), and have been elaborated extensively in numerous studies (Hawkins et al., 2009; Xu, Wickramarathne, & Chawla, 2016; Cui et al., 2016; Persson, Bohlin, Edler, & Rosvall, 2016). Clones are created by identifying the states in a lower-order model that need to be split, and then relearning the counts. A simple and effective way to decide how to allocate clones to each symbol is to do so proportionally to the number of times that the symbol appears in the data. The intuition that more frequent symbols need more clones is obvious when we go to the extreme case of memorizing a sequence: we need exactly as many states as total symbols appear in the sequence, and as many clones per symbol as occurrences of that symbol appear in the sequence.

### Training CHMMs with memorize-generalize

In (Dedieu et al., 2019) we show that a CHMM can be learned using EM. Briefly, EM starts from a random connectivity pattern and smoothly improves the model to fit it to the data. This method is guaranteed to converge to a local maximum of the likelihood. The restricted capacity of the model (number of clones) prevents the CHMM from memorizing the data, thus resulting in a model that generalizes to new, unseen sequences.

Despite its advantages, the EM algorithm is not very biologically plausible. Instead, we developed a memorize-generalize algorithm for training CHMMs. First, the CHMM is forced to memorize a *training sequence* (see Fig. 1B). At this point, the CHMM can only generate or recognize verbatim portions of the training sequence. Then the CHMM is endowed with a *random jump* probability using a standard pseudocount in its transition matrix (Fig. 1C), which introduces a small probability of jumping to any arbitrary clone at each time instant. Finally, a *generalization sequence* is presented. In an online manner, the CHMM decodes^1^ the next symbol according to its current model, and the newly discovered transition is added to a count matrix. The transition matrix of the CHMM is immediately updated with a normalized version of the count matrix, and then the next symbol is decoded. This online adaptation is iterated. Note that the training sequence can also be used as generalization sequence, and that presenting it multiple times (or *epochs*) will continue to modify the model.

The memorize-generalize algorithm can be interpreted as an online form of Viterbi training (Jelinek, 1976). The immediate acquisition of data (memorization) and the sequential distillation process involving only temporally local data (generalization) makes memorize-generalize more biologically plausible than a global and slow learning algorithm like EM, even if its performance is oftentimes inferior.

## Modeling long-range structure in bird songs

### Bird song as a model of sequence learning

The avian song system has provided a rich system for understanding how complex sequential behaviors are produced by the brain, including how they are learned through observation and practice (Mooney, 2014). Similar to human speech, avian song is built from elementary units known as syllables (Doupe & Kuhl, 2002). Each syllable is produced by a different sequence of action potential bursts in the premotor cortical area, distinct syllable types are produced by largely non-overlapping neural sequences (Long, Jin, & Fee, 2010). These syllables are repeated in groups to form phases, which are eventually sequenced to form songs. It has been proposed that the link between the neural sequences corresponding to different syllables is mediated by a feedback loop through the thalamus and as such songs are generated through a distributed circuit that spans the avian forebrain and thalamus (Hamaguchi, Tanaka, & Mooney, 2016). It is important to note that sequences in non-human primate brain are also generated by a similar distributed circuit (Sommer & Wurtz, 2008).

During sensory learning, the juvenile songbird listens to and memorizes the song of a tutor (Mooney, 2014; Doupe & Kuhl, 2002). These juveniles initially produce a highly variable *subsong* in which the syllable order often gets mixed up, and eventually arrange the newly-learned syllables into the correct order (Liu, Gardner, & Nottebohm, 2004; Lipkind et al., 2017). Notably, sensory learning is remarkably fast as tutor song can be accurately mimicked only after a few (several hundred) repetitions (Mooney, 2014). Song learning is associated with a gradual change in neural sequence structure (Okubo, Mackevicius, Payne, Lynch, & Fee, 2015) — with a lack of structure during the subsong stage (protosequences), followed by an increase in rhythmic bursting duration as the bird acquires more syllables. A recent study proposed that new syllable types can emerge by the gradual splitting of a single protosequence (Okubo et al., 2015). During the splitting process, Okubo and colleagues observed neurons specific to each of the emerging syllables, as well as shared neurons that were active in both syllable types. Splitting a neural sequence in this fashion is computationally efficient as it allows learned components of a primitive motor program to be reused in each of the daughter sequences. This is akin to the way in which memorize-generalize reuses previously memorized subsequences during the generalization phase.

## Results

By analyzing temporal correlations between syllables within a song, several studies have noted that the choice of what to sing next is determined not only by the current syllable but also by previous syllables sung (Markowitz, Ivie, Kligler, & Gardner, 2013; Katahira, Suzuki, Okanoya, & Okada, 2011). This means that syllable choices in a particular context depend on the history of the song. Therefore, uncovering the statistics of these long-range correlations may provide a crucial insight into how the brain assembles complex behaviors from primitives in a context dependent manner.

To illustrate the efficacy of our CHMM in learning higher-order temporal structure, we analyzed songs from six canaries (Markowitz et al., 2013). Briefly, as described by Markowitz and colleagues, spectrograms of the songs from six canaries (*Belgian Waterslager*) were manually annotated and identified syllables were converted to strings. For each bird, we first created 30 different 90%-10% cross-validation (CV) train-test splits. Next, for each split we ran both the memorize-generalize and the EM algorithm for multiple epochs and measured the negative log-likelihood in both the training and test sets. These results, averaged over CV splits, are shown in Fig. 2A. Note that a lower negative log-likelihood indicates better performance of the model. Notably, we found only a marginal difference when using lookahead buffers of different sizes (dashed vs. solid line in Fig. 2A). Although training with EM produces faster convergence than memorize-generalize (bottom panel Fig. 2A), our methods outperform the one used by Markowitz and colleagues on all six canaries (three of them shown in Fig. 2, see Fig 6 in Markowitz et al.). Given that the transition matrix of the CHMM is a higher-order graph, we are able to visualize the relationships between syllables in the song (Fig. 2B), which led to two major observations. First, transitions between some syllables were more probable than others, suggesting a fine-grained ordering of syllables, which would likely have some ethological meaning among conspecifics. Second, syllables within a song have long-range temporal relationships. Together, these observations support the notion of contextual dependencies within a song (Katahira et al., 2011). Relating these contextual-dependencies between syllables with neurophysiological properties of the canary brain will not only provide us with an understanding of the neural circuits involved in song generation, but will also reveal how the brain uses contextual information during speech (Doupe & Kuhl, 2002).

**Figure 2:**
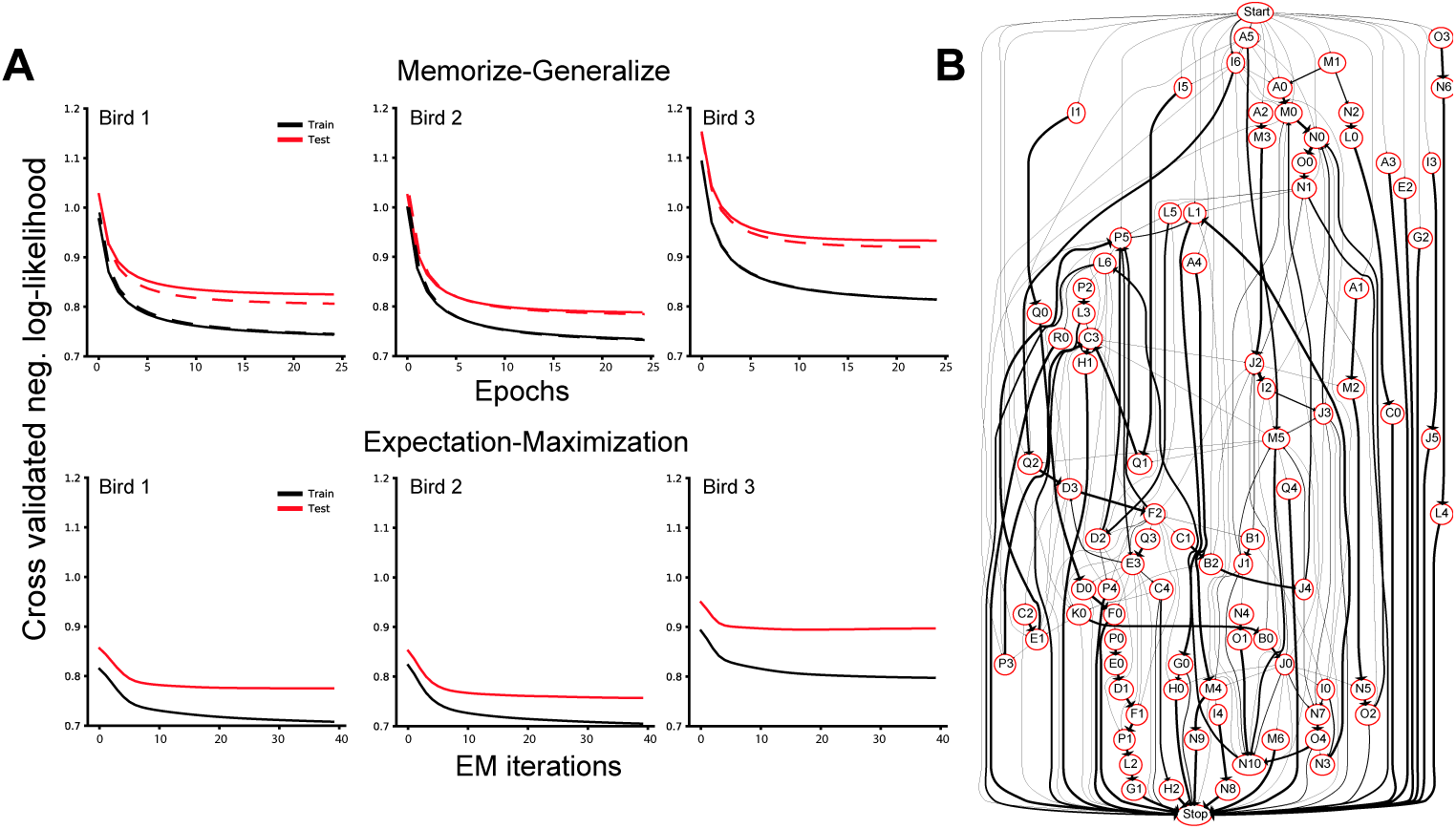
CHMM trained with memorize-generalize algorithm uncovers long-range dependencies between syllables in canary song. (A) Comparison between memorize-generalize (top) and expectation-maximization (bottom) for three representative birds. In the top panel, dashed lines are the memorize-generalize algorithm with a lookahead buffer *B* = 2, while the solid line is a greedy version of the algorithm (*B* = 0). (B) Visualization of the transition matrix for an example bird. Thicker edges represent higher transition probability. Each observed syllable is identified with a different letter, whereas each clone is represented with a different number.

## Conclusions

In this paper, we drew inspiration from sequence representation in the cortex to develop a new sequence learning model — the CHMM. When trained with a new online learning algorithm, the CHMM was efficient at learning higher-order temporal correlations between syllables in birdsong. Unlike other state-of-the-art sequence learning models, such as LTSMs or n-grams, our learned model was effective at dealing with uncertain contexts, which is a crucial requirement for agents operating in the real world.

## Footnotes

1 Viterbi decoding assumes that the clones decoded so far are known and fixed, and a small lookahead buffer for the next *B* symbols is available.

